# The effects of postural threat induced by a virtual environment on performance of a walking balance task

**DOI:** 10.1101/752139

**Authors:** Amir Boroomand-Tehrani, Andrew H. Huntley, David Jagroop, Jennifer L. Campos, Kara K. Patterson, Luc Tremblay, Avril Mansfield

## Abstract

Rapid motor learning may occur in situations where individuals perceive a threat of injury if they do not perform a task well. This rapid motor learning may be facilitated by improved motor performance and, consequently, more errorless practice. As a first step towards understanding the role of perceived threat on rapid motor learning, the purpose of this study was to determine how performance of a motor task is affected in situations where perceived threat of injury is high. We hypothesized that perceived threat of injury in a virtual environment would result in improved performance of a walking task (i.e., walking on a narrow beam). Results demonstrated that increased perceived threat of injury yielded slightly greater, but not statistically significant, balance performance in virtual environments (median percentage of successful steps: 78.8%, 48.3%, and 55.2% in the real low-threat, virtual low-threat, and virtual high-threat environments, respectively). These results may be partially attributed to habituation to threat over time and practice. If implemented carefully, virtual reality technology can be an effective tool for investigating walking balance in environments that are perceived as threatening.

## INTRODUCTION

There is evidence that rapid motor learning can occur in situations where errors can yield an injury. For example, fall-resisting skills can be rapidly acquired through repeated exposure to slips during sit-to-stand tasks (Pavol, Runtz, Edwards, & Pai, 2002; Pavol, Runtz, & Pai, 2004) and walking balance tasks, such as preventing a backwards loss of balance (Bhatt, Wening, & Pai, 2006; Bhatt, Yang, & Pai, 2012; Pai, Yang, Bhatt, & Wang, 2014). Individuals can adapt to these perturbations within just a single session by rapidly improving both proactive and reactive balance control strategies, such as better centre of mass (COM) stability and reactive stepping (Bhatt et al., 2006; Pai, Bhatt, Yang, & Wang, 2014; Pai, Yang, et al., 2014). These fall-resisting skills are often retained beyond the single session in which they were acquired (e.g., up to six months later; Pai, Bhatt, et al., 2014; Pai, Yang, et al., 2014). Losses of balance often lead to falls that can ultimately cause injuries. The perceived potential for injury associated with a loss of balance may further trigger central nervous system (CNS) activity to quickly learn fall-resisting skills, which can be retained for an extended period (Adkin, Frank, Carpenter, & Peysar, 2000; Pai, Bhatt, et al., 2014). Furthermore, rapid balance skill acquisition is particularly evident among individuals who are more at-risk of falls, as the fear developed from previous falls might further motivate the rapid adaptation (Pai, Bhatt, et al., 2014).

Adaptive responses to threat mobilize energy to the heart, muscles, and brain to carry out the necessary actions for surviving the threatening situation (Sapolsky, 2000), which may involve dedicating resources for optimizing motor performance. We suggest that rapid motor learning in these threatening situations may be facilitated by enhanced motor performance and consequently, more errorless practice. Errorless motor learning is a form of implicit learning which involves reducing the number of outcome errors during practice (Capio, Poolton, Sit, Eguia, & Masters, 2013). The rapid motor learning of balance skills in situations of high perceived threat may be facilitated by errorless practice, as individuals resist errors to avoid injury. However, the role of perceived threat of injury on rapid motor learning is still unclear and requires further investigation.

Immersive virtual reality (VR) technology is used to simulate a realistic virtual environment (VE) by providing the user with sensory information that resembles objects and events in the real world (Slater, 2009). VEs can be used to evaluate motor performance in environments that would otherwise be avoided due to feelings of fear and a lack of safety (Cleworth, Horslen, & Carpenter, 2012). VR technology can be a particularly useful tool for studying the effects of perceived threat on balance, as it is possible to control and customize threatening stimuli without compromising safety.

A previous study did not find any significant differences in balance performance when comparing beam walking in VEs with and without threatening heights (Peterson, Furuichi, & Ferris, 2018). This previous study included a secondary cognitive task, which may have diverted attentional resources away from the balance task, potentially interfering with the dedication of resources to optimize motor functioning in the high-threat condition. As a first step towards understanding the role of perceived threat on rapid motor learning, the purpose of the current study was to determine if performance of a motor task improves in situations where perceived threat of injury is high. We were specifically interested in the influence of context-specific threat of injury; that is, the introduction of a scenario for which participants would perceive that an injury may occur if they did not perform the task well. We hypothesized that perceived threat of injury, due specifically to a loss of balance, in a VE would result in improved motor performance in a balance beam task. This work was also intended to inform future research by providing insight on the effectiveness of using VR technology to study threat and walking balance.

## METHODS

### Participants

Twenty-four participants (12 males) with a mean age of 22.8 years (*SD* = 1.8 years) completed the study. Participants were excluded if they had: difficulty understanding verbal or written English; history of epilepsy, or other neurological conditions that could affect balance or mobility or prevent use of VR hardware; history of motion sickness; diagnosed acrophobia; recent injury to the eyes, face, or neck that would prevent comfortable use of VR hardware; surgery or recent injury to the lower extremity that could affect balance or mobility; difficulty in hearing that was not corrected with a hearing aid; difficulty in vision that was not corrected with contact lenses; sensitivity to flashing light or motion; and/or 6 months of formal dance or gymnastic training in the last 10 years. Due to hardware limitations, glasses were not permitted and contact lenses were required for those who usually require corrective lenses during daily activities. Participants were also excluded if they had participated in previous studies that involved walking on a balance beam, as prior experience may affect the results. Written informed consent was obtained from all participants. The University Health Network Research Ethics Board approved the study protocol and all other study materials. Participants received a $30 gift card as compensation.

### Apparatus

Participants completed a balance beam-walking task in 3 separate conditions, and the sequences of conditions were counterbalanced across participants. The balance beam used in this study was 8.5 cm tall, 300 cm long, and 3.8 cm wide. Figure 1A shows the testing area where participants completed the balance task.

**Figure 1:**
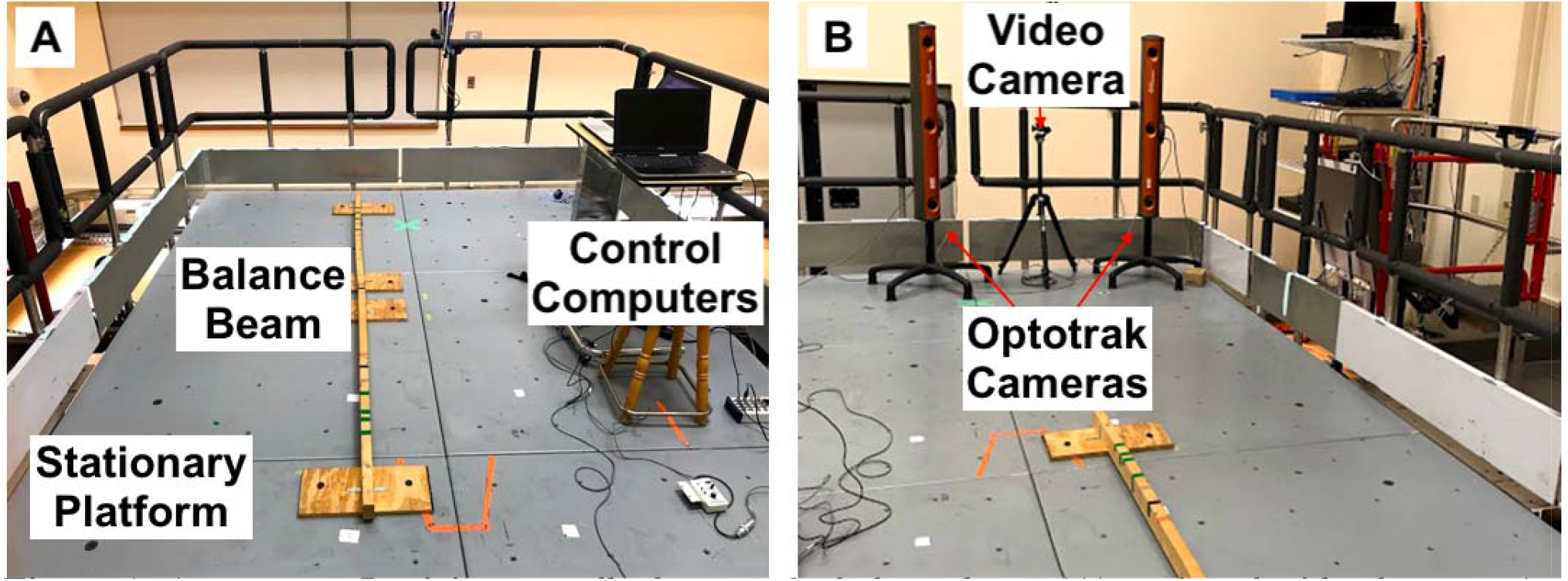
Apparatus. Participants walked across the balance beam (A) equipped with a harness. An experimenter walked along each side of them for added safety precautions. The cameras (B) were positioned behind the participant to record marker position and video footage during beam passes.

### Data collection

Two Optotrak 3D Investigator (Northern Digital Inc., Waterloo, Canada) cameras were used to measure kinematic data, as well as to update the visual scene in the VE in real time (Figure 1B). The Optotrak cameras tracked the three-dimensional positions of markers placed at the neck, as well as markers placed on the head to update the visual scene. Motion data were recorded at 100 Hz. Data collection sessions were also recorded using a digital video camera.

Electrodermal activity (EDA) data were collected throughout the beam walk by placing electrodes on the palmar surfaces of the index and middle fingers of the non-dominant hand. Reusable silver/silver chloride finger electrodes filled with conductive paste were placed on the fingers. EDA was sampled at 100 Hz. The Endler Multidimensional Anxiety Scale – Trait (EMAS-T; Endler, Edwards, Vitelli, & Parker, 1989) was administered at the beginning of each data collection session to evaluate trait anxiety. The Endler Multidimensional Anxiety Scale – State (EMAS-S; Endler et al., 1989) was administered after completing the task in each condition to evaluate participants’ perceptions of the preceding task. This scale allowed us to compare situation-induced anxiety to baseline levels of general anxiety. The International Physical Activity Questionnaire (Craig et al., 2003) was administered at the beginning of the experiment to evaluate physical activity over the previous week.

### Virtual reality system

An Oculus DK1 (Oculus VR, Menlo Park, California, USA) head mounted display (HMD) was used to immerse the participant in the VE. The HMD was 1280 pixels wide x 800 pixels high (640 x 800 per eye) with a 110º field of view and displayed stereoscopic graphics. The VE contained visual depth cues, resembling depth cues in the physical lab, that helped participants perceive the VE in 3 dimensions. The viewpoint in the VE was controlled by motion capture data from the Optotrak cameras, which synchronized the participants’ position and movement using a rigid body with infrared light emitting diodes attached to the HMD. By determining head and body position, the Optotrak cameras updated the visual scene of the HMD accordingly; this allowed the viewpoint to update in real-time so that the visual motion was consistent with the physical motion during walking. Use of the HMD ensured that participants did not gain any visual inputs from the real world.

### Protocol

Participants started with their non-dominant foot on the beam and were instructed to attempt to walk across the entire balance beam without stepping off, and if they stepped off the beam, to step back on and resume walking at the point of descent. There were no specific instructions regarding how to control their upper limbs. Following completion of a beam pass, participants returned to the starting position for subsequent trials. Participants completed the balance beam walking task in a block of 4 trials for familiarization, and blocks of 10 trials in each of the 3 testing conditions: 1) low-threat real environment; 2) low-threat VE; and 3) high-threat VE (see below for more details about these environments).

#### Familiarization trials

Participants first performed 4 trials, 2 in a low-threat real environment and 2 in a low-threat VE, to familiarize themselves with the task. At the beginning of every trial in a VE, participants completed a series of object identification tasks as part of an immersion protocol (Figure 2A). Specifically, they verbally identified the shape and colour of all objects and answered questions regarding their proximity. For example, they were asked to state the colour of the nearest cube or furthest sphere. Successful completion of the object identification tasks confirmed that participants were able to perceive relative depth within the VE. They were also instructed to contact the beam in various ways, such as by tapping and stepping on it, to understand that the virtual beam was matched with the physical beam in space. This immersion period also allowed participants to acclimate to the VE and adapt to mechanical factors, such as the weight of the HMD, to ultimately enhance their sense of presence (Cleworth et al., 2012). This protocol was not intended to test participants’ ability to make relative depth judgments in the VE.

**Figure 2:**
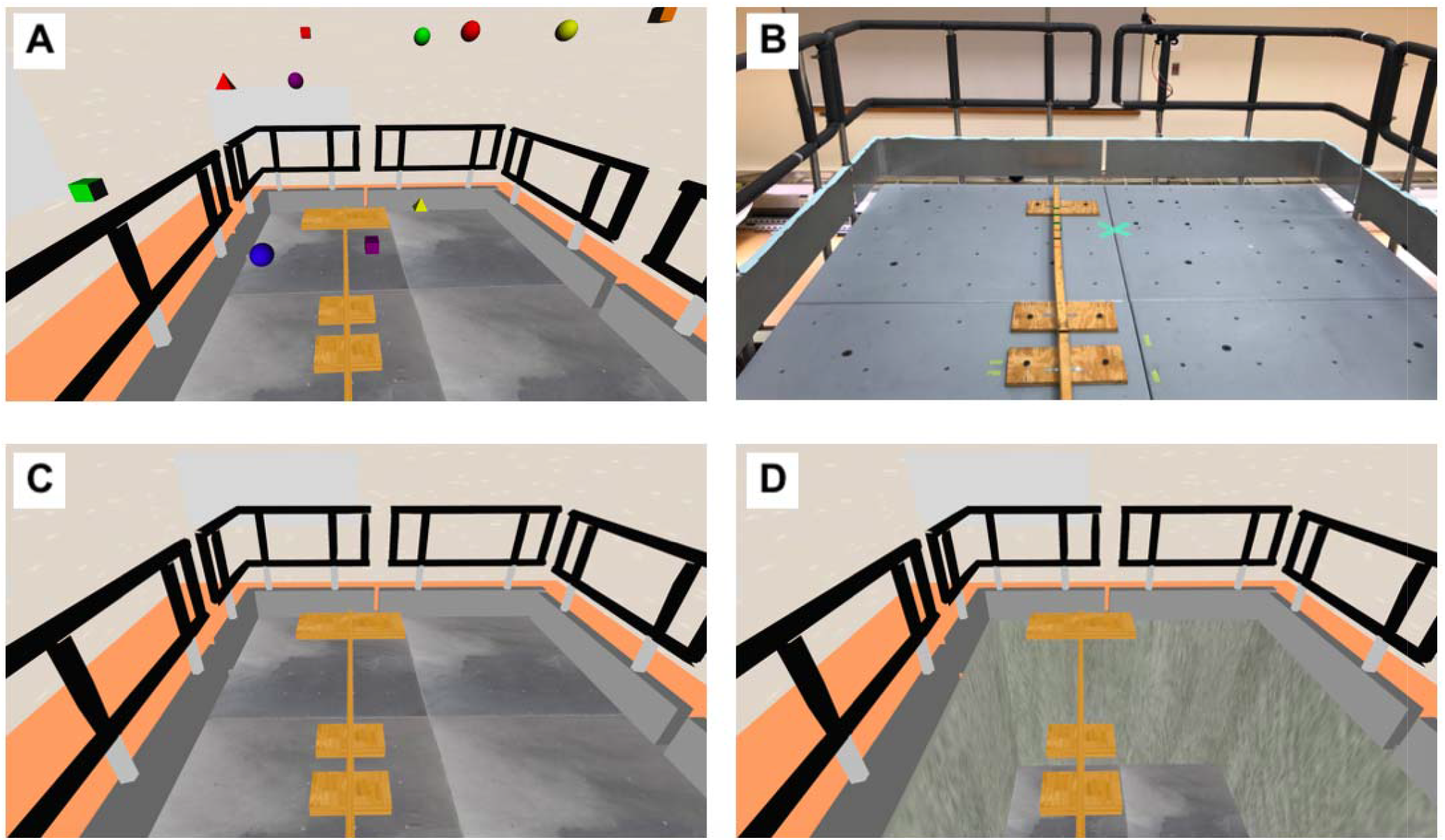
Environments. Before walking on the beam in a VE, participants completed an immersion protocol wherein they answered questions regarding the relative positions of virtual objects in the VE (A). While walking in the low-threat real environment (B), participants were equipped with VR hardware equalize mechanical factors between conditions. The low-threat VE resembled the real environment (C). The high-threat VE was identical to the low-threat VE, except that the virtual platform descended at the beginning of each trial (D).

#### Low threat, real environment (Figure 2B)

While walking in the real environment, participants wore the HMD over their forehead (i.e., not obscuring their eyes) to adapt to mechanical factors, such as the added weight, which may affect postural control. Participants also wore basketball ‘dribble’ goggles that obscured the lower portion of their field of view, to approximate the field of view of the visual display of the HMD. This hardware was worn to resemble the experience of walking in a VE, as gait parameters can differ due to the weight of the HMD and smaller field of view (Mohler, Campos, Weyel, & Bulthoff, 2007).

#### Low threat, VE (Figure 2C)

The VE in the low-threat condition simulated the real environment. A virtual rendering of the beam was visible in the VE in the ‘real world’ location of the beam. As the virtual beam and real beam were matched in space, a step off the real beam resulted in a step off the virtual beam. There were no visual consequences to stepping off the beam in this VE.

#### High threat, VE (Figure 2D)

In the high-threat VE condition we virtually simulated the balance beam to be elevated by 10 m. This height appeared to be dangerous to participants and has been previously shown to be sufficient to produce feelings of threat (Cleworth et al., 2012). A step off the beam resulted in bright red flashes representing a negative consequence.

### Data processing

The primary outcome was performance of the balance beam-walking task, quantified by step success, which was determined from the video footage. A step was defined as a success when the swing foot landed and stabilized on the beam, such that they were able to attempt the next step. A step was defined as a fail when the swing foot contacted the floor, or contacted the beam but then immediately contacted to the floor. A step was also considered a fail if the stance foot contacted the floor as the swing foot approached the beam. Performance was measured by counting steps on the beam, as a proportion of total steps taken (i.e., percentage of ‘successful steps’) and the numbers of steps taken before the first failed step. The time to first fail was calculated as the time elapsed between the onset of the first attempted step and first failed step (estimated from video footage).

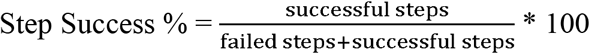

Secondary outcomes were EDA, movement variability, and state anxiety. EDA data were low-pass filtered at 5 Hz using a 2^nd^ order Butterworth filter. To separate the influence of physical activity on physiological arousal and measure physiological stress as a function of walking condition, we normalized EDA by recording EDA during each beam walk and subtracting EDA values from a one-second standing baseline immediately before the beam walk.

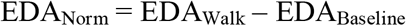

Electrodermal responses can occur up to 5 seconds following a stimulus (Benedek & Kaernbach, 2010). Therefore, EDA values that were recorded up to 5 seconds after completion of a walking trial were included as part of the beam walk.

Kinematic data were filtered using a zero-phase-lag low-pass 4^th^ order Butterworth filter with a cut-off frequency of 6 Hz. Movement variability was calculated as the standard deviation of the medial-lateral positions of the markers on the neck, only during instances when the participant was walking on the beam (Domingo & Ferris, 2009), using Visual3D (C-Motion, Germantown, Maryland, USA).

### Data analysis

Mean percentage of successful steps, EDA_Norm_, and movement variability were calculated for each participant and each condition prior to analysis. Non-parametric Friedman tests were used to compare successful steps, EMAS-state, number of successful steps before first failure, and movement variability between conditions. When a significant difference was observed between the three conditions, a Wilcoxon signed-rank test was used for further pairwise comparisons between the: 1) low-threat real environment and low-threat VE; and 2) low-threat VE and high-threat VE. A Bonferroni-adjusted significance level was calculated for post-hoc testing to account for the increased likelihood of a Type I error when making multiple comparisons (adjusted alpha=0.025). The two low-threat environments were compared to evaluate the effects of using VR, and the two VEs were compared to evaluate the effects of perceived threat. All statistical tests were performed in SAS (Version 9.4, Cary, North Carolina, USA). EDA_Norm_ was log transformed to fit a normal distribution. A one-way repeated measures analysis of variance (ANOVA) was used to compare transformed EDA_Norm_ between the three conditions. When a difference was observed between the three conditions, a Tukey’s honestly significant difference post hoc test was used for further pairwise comparisons between the: 1) low-threat real environment and low-threat VE; and 2) low-threat VE and high-threat VE.

It is possible that feelings of perceived threat of injury may subside with repeated exposure to the virtual threat, particularly if participants commit errors without the expected consequences in VEs (e.g. by stepping off the beam but not falling to the floor). Thus, habituation to the threat, measured by EDA, was investigated to reveal any changes in perceived threat.

## RESULTS

Twenty-seven participants were recruited; however, three did not complete the full protocol due to feeling motion sick. Therefore, 24 participants completed the full protocol and were included in the analysis (Table 1).

**Table 1:** Participant characteristics. Values shown are mean ± standard deviation. The maximum possible score for each EMAS-T subscale is 75.

|  |  |
| --- | --- |
| Age (years) | 22.8 ± 1.8 |
| Height (m) | 1.7 ± 0.1 |
| Weight (kg) | 70.7 ± 15.5 |
| Sex (number) |  |
| Male | 12 |
| Female | 12 |
| EMAS-T subscales (score) |  |
| Social Evaluation | 40 ± 8.0 |
| Physical Danger | 57 ± 8.4 |
| Ambiguous Situations | 42 ± 11.6 |
| Daily Routines | 23 ± 5.3 |
| IPAQ (metabolic equivalent of task minutes) | 4632 ± 4265.9 |
| EMAS-T=Endler Multi-dimensional Anxiety Scale – Trait (Endler et al., 1989) |  |
| IPAQ=International Physical Activity Questionnaire (Craig et al., 2003) |  |

### Step success

Balance beam-walking performance was significantly different between the 3 conditions (χ^2^(2) = 43.80, χ p < 0.0001; Table 2). Although step success appeared to be higher in the high-threat VE compared to the low-threat VE, this difference was not statistically significant (S = 57.5, p = 0.10; Table 2). The number of successful steps before first fail was significantly different between the 3 conditions (χ^2^(2) = 36.77, p < 0.0001) with a greater number of steps in the low-threat real environment compared to the low-threat VE (S = 150, p < 0.0001). Time to first fail was significantly different between the 3 conditions (χ2(2) = 27.41, p < 0.0001) with a longer time to fail in the low-threat real environment compared to the low-threat VE (S = 127, p < 0.0001). There were no significant differences in movement variability between conditions (χ^2^(2) = 4.58, p = 0.10; Table 2).

**Table 2:** Outcome measures between conditions. Values are presented as medians (interquartile ranges) for all balance and state anxiety outcomes, and their p-values are from the Friedman tests comparing outcome measures between conditions. Values are presented as means (standard deviation) for all EDA outcomes, and their p-values are from the repeated measures ANOVA comparing the transformed EDA values between conditions. For post-hoc testing, values are from the Wilcoxon signed-rank test for pairwise comparisons of balance and state anxiety outcomes. The Bonferroni-adjusted significance level is α=0.025. The maximum possible score for the EMAS-S is 100.

|  | Real +<br>Low<br>Threat<br>(RL) | Virtual +<br>Low<br>Threat<br>(VL) | Virtual +<br>High<br>Threat<br>(VH) | *p-value*<br>(Friedman<br>test) | *p-value*<br>(RL vs.<br>VL) | *p-value*<br>(VL vs.<br>VH) |
| --- | --- | --- | --- | --- | --- | --- |
| Step success (%) | 78.8<br>(20.6) | 48.3 (11.7) | 55.2 (17.8) | <0.0001 | <0.0001 | 0.10 |
| Steps before first<br>fail (number) | 2.8 (3.6) | 0.9 (0.4) | 1.0 (0.6) | <0.0001 | <0.0001 | 0.060 |
| Time to first fail<br>(s) | 2.7 (1.9) | 1.1 (0.7) | 1.4 (0.9) | <0.0001 | <0.0001 | 0.037 |
| Movement<br>variability (cm) | 4.62<br>(1.98) | 2.89 (0.94) | 4.04 (2.56) | 0.10 | - | - |
| EMAS-S score | 36 (19.5) | 46 (20) | 53 (20.5) | 0.0042 | 0.0026 | 0.0010 |
| EDA<sub>Norm</sub> (μS) | 0.23<br>(0.16) | 0.23 (0.12) | 0.24 (0.12) | 0.88 | - | - |

EMAS-S=Endler Multidimensional Anxiety Scale – State (Endler et al., 1989)
EDA=electrodermal activity

### Anxiety

There were no significant differences in EDA_Norm_ between conditions (F(2,69) = 0.13, p = 0.88; Table 2). Changes in EDA_Norm_ were analyzed over the course of all 10 trials within a block to explore any trends pertaining to habituation to threat (Figure 3). EDA_Norm_ decreased over the course of a beam walking block for all 3 conditions. We measured changes in EDA_Norm_ from trials 1 to 2 and 2 to 10 to determine how much of the habituation was attributed to the first trial. In the real low-threat condition, EDA_Norm_ decreased by 0.06 µS and 0.04 µS from trials 1 to 2 and 2 to 10, respectively. In the virtual low-threat condition, EDA_Norm_ decreased by 0.05 µS and 0.09 µS from trials 1 to 2 and 2 to 10, respectively. There was a considerable drop in EDA_Norm_ after just 1 trial in the high-threat condition, as EDA_Norm_ decreased by 0.14 µS and 0.03 µS from trials 1 to 2 and 2 to 10, respectively.

**Figure 3:**
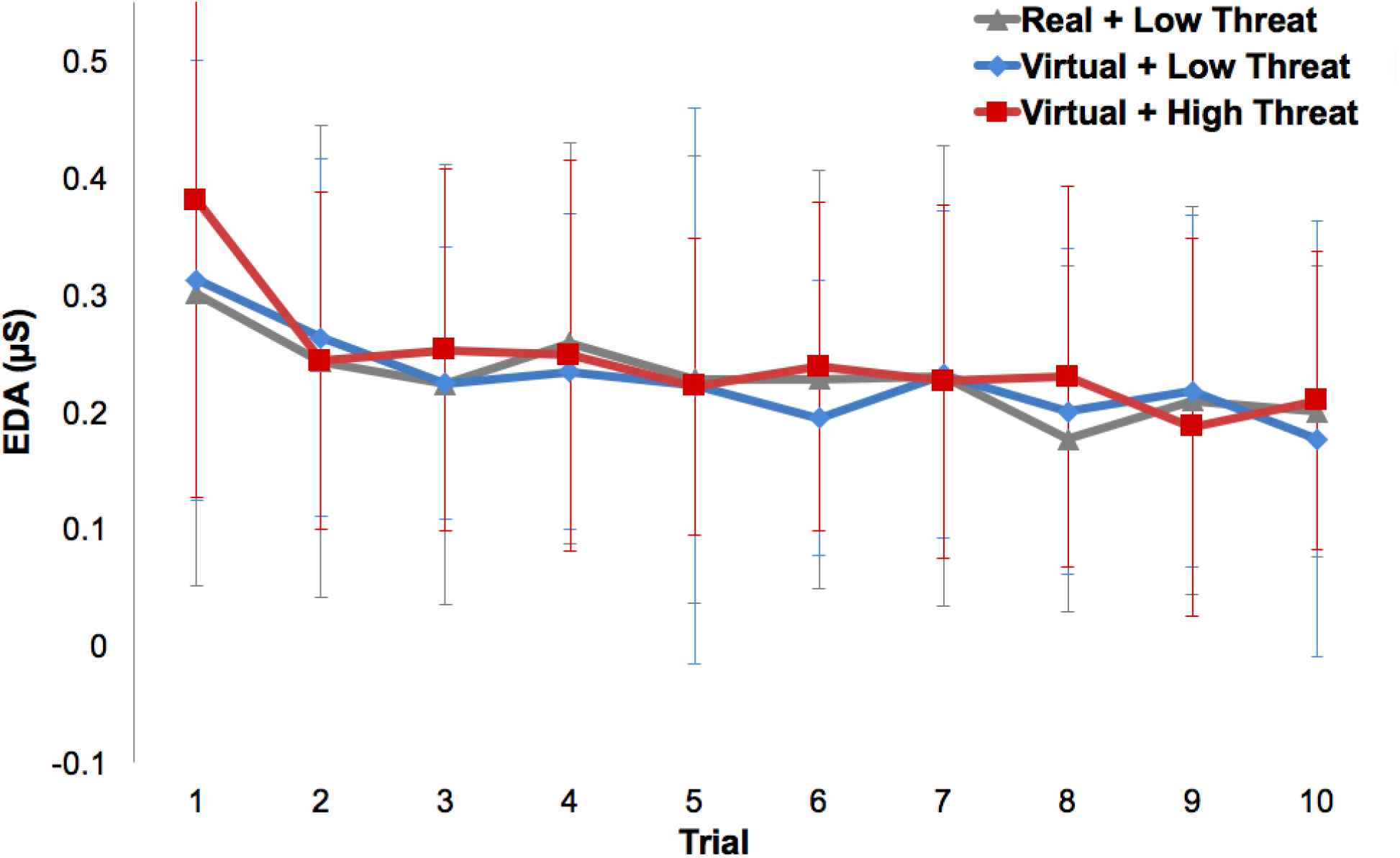
EDA changes throughout trials. Mean EDA_Norm_ is shown for each walking trial. The standard deviation for each trial and condition is displayed as horizontal bars.

EMAS-S scores differed significantly between conditions (χ^2^(2) = 3.51, p = 0.0017; Table 2). EMAS-S scores were significantly higher in the high-threat VE compared to the low-threat VE (S = 99, p = 0.0010; Table 2).

## DISCUSSION

The purpose of this study was to determine if performance of a balance beam walking task improves in situations where perceived threat of injury is high. We hypothesized that performance would be better in situations where an individual perceived a threat of injury associated with not performing the task well. We used a VE to produce perceived threat. While the increased perceived threat yielded slightly greater step success in the high-threat VE compared to the low-threat VE, the difference was not statistically significant. This finding did not corroborate with previous studies that demonstrated statistically significant improvements in task performance when individuals perceived a threat of injury if they did not perform well in cognitive and simple motor tasks (Abend et al., 2013; Adkin et al., 2000; Bhatt et al., 2006; Bhatt et al., 2012; Brown & Frank, 1997; Collins, Mendelsohn, Cain, & Schiller, 2014; Pai, Bhatt, Wang, Espy, & Pavol, 2010). Similarly, in the current study there was no statistically significant difference in movement variability between the three conditions. We predicted that movement variability would be lower in the high-threat condition than the low-threat condition due to improved balance control. Surprisingly, higher movement variability coincided with greater step success across the three walking environments, which contradicts our expectations. Although it has been reported that individuals engage in tighter control of posture under conditions of increased postural threat (Adkin et al., 2000; Brown & Frank, 1997; Carpenter, Frank, & Silcher, 1999), these findings were not reproduced in this study. We only calculated movement variability for instances when participants were stable on the beam. Assessing movement variability under these limited circumstances may have contributed to the differences with previous findings. When nearing a loss of balance in the low-threat VE, participants may have conceded the fail and stepped off the beam. However, participants in the high-threat VE were likely more motivated to not concede fails as easily. Instead, in the high-threat VE condition they may have been attempting more reactive trunk adjustments to control their centre of mass, resulting in more trunk variability, in order to not lose their balance on the beam. Another perspective suggests that increased movement variability reflects exploration of an efficient movement pattern in a changing environment (Bartlett, Wheat, & Robins, 2007; Jarvis, Smith, & Kulig, 2014). Exploratory behaviours may have been prompted as the beam walking task was a novel experience for all participants. Although time to first fail was greater in the high-threat VE than in the low-threat VE, this difference was not statistically significant. Time to first fail may reflect increased motivation to maintain balance when perceiving a threat.

When comparing the two low-threat environments (real and virtual), step success, the number of successful steps before the first fail, and time to first fail were significantly greater in the real environment than in the VE. This mismatch in step success may indicate a need for technical improvements or reflect inherent differences in walking in a VE, such as having fewer sensory cues; it is likely a combination of both factors. Posture has previously been reported as less stable in VEs (Janeh et al., 2017). Any conflict between the visual, somatosensory, and vestibular information used to control balance can negatively affect balance performance (Fitzpatrick & McCloskey, 1994; Kelly, Riecke, Loomis, & Beall, 2008; Nishiike et al., 2013; Peterson et al., 2018). End-to-end latencies in a VE reflect the time elapsed between head movement and the consequent updating of the visual scene. These latencies may cause conflict between the visual and vestibular systems. The HMD used in this study had an end-to-end latency of 50-60ms, and although this level of latency is not an issue for static tasks, large head movements are often not well translated by HMDs (Robert, Ballaz, & Lemay, 2016). Sensory conflict, especially in dynamic balance tasks where visual information is important, could negatively affect walking performance (Robert et al., 2016). The significant discrepancy between real and virtual environments supports our rationale for comparing virtual experiences against each other to evaluate the effects of perceived threat.

Increasing perceived body elevation in a VE has been previously found to increase feelings of perceived threat in both static and dynamic balance tasks (Cleworth et al., 2012; Ehgoetz Martens, Ellard, & Almeida, 2015; Peterson et al., 2018). The current study reported similar findings, as the high-threat VE produced significantly greater state anxiety than the low-threat VE. Continuing to improve the technical specifications of the VE would further contribute to a more convincing experience, and as a result a more convincing threat. Although there were no significant differences in mean EDA between the three conditions, the trial-by-trial changes in EDA revealed noteworthy trends presumed to be associated with habituation to threat. EDA declined substantially following the first trial in all three conditions. This decline continued until the completion of all 10 trials. The reduction in EDA from the first to second trial was most apparent in the high-threat VE, as the initial illusion of threat may have been lost following participants’ first instances of failure without actual negative consequences. The trial-by-trial waning of EDA may have also affected self-reported state anxiety. The EMAS-S was administered at the end of the trial block, when EDA had already declined substantially. The initial anxiety from the high-threat condition may have been depleted by the time the questionnaire was completed. If individuals habituated to feelings of perceived threat, there is likely a downstream effect on the performance measures, such as step success and movement variability, that we predicted to improve as a result of perceived threat.

## CONCLUSIONS

Although step success was slightly greater in the high-threat VE than in the low-threat VE, this difference was not statistically significant. There is evidence that participants did feel more threatened in the high-threat VE. However, the manipulation to produce feelings of perceived threat may not have been sufficiently robust to significantly affect performance. Decreased anxiety was likely partially attributed to participants’ waning perception of threat as they realized the threat was not real when they committed their first fails. Habituation to threat likely affected self-reported state anxiety and walking performance, both of which we predicted to increase in the high-threat condition. More robust manipulations should be tested to provoke the adaptive reactions that facilitate enhanced motor control. To further understand the role of perceived threat in promoting rapid motor learning, future studies should investigate how varying levels of perceived threat, committed errors, and difficulty of task, affect motor learning.

## Acknowledgements

This work was supported by the Natural Sciences and Engineering Research Council of Canada (RGPIN-2014-04199) and the Ministry of Research and Innovation (Ontario). The authors acknowledge the support of the Toronto Rehabilitation Institute; equipment and space have been funded with grants from the Canada Foundation for Innovation, Ontario Innovation Trust, and the Ministry of Research and Innovation. Andrew Huntley was supported by an award from Heart and Stroke Foundation Canadian Partnership for Stroke Recovery. Avril Mansfield is supported by a New Investigator Award from the Canadian Institutes of Health Research (MSH-141983). Jennifer Campos is supported by a Canada Research Chair (Tier 2). These funding sources had no role in the design or execution of this study, analyses or interpretation of the data, or decision to submit results. We would also like to thank Suluxshiga Jeyendran, Rebecca Graham, Nastasia Kujbid, and Tong Yu for their assistance with data collection and processing.

